# Genome-wide association study and sequence similarity analysis for unilateral renal agenesis using heterogeneous stock rats undercovers the *KIT* gene and *AHR*, *ATF3*, *GATA3*, *HNF1B*, *POU2F2*, and *TFCP2* transcription factors as potential candidates to explain incomplete penetrance

**DOI:** 10.1101/2023.09.19.558496

**Authors:** Joel D. Leal-Gutiérrez, Daniel Munro, Denghui Chen, Riyan Cheng, Tengfei Wang, Hao Chen, Paul Meyer, Keita Ishiwari, Terry E. Robinson, Christoph Rau, Michael Garrett

## Abstract

Human unilateral renal agenesis is a congenital urinary tract malformation. Affected individuals have only one kidney, which is often an asymptomatic developmental defect. A total of 5,585 male and female HS rats were assessed for unilateral renal agenesis and genotyped for 3’513,321 markers. The R package SAIGEgds was used for the association analysis. The adjusted p-value threshold for the association analysis determined by permutation was equal to 5.6 (-log10). Two additional datasets were used as validation tests. Population two included 1,577 rats genotyped for 7,425,889 markers and a case-control imbalance equal to 1:174; population three included 1,407 rats, genotyped for 254,932 markers and case-control ratio equal to 1:38. The python package GxTheta was used to perform a polygenic epistasis analysis for the analyzed HS rat population. A founder haplotype mosaic determination was performed using the R package QTL2. Associated regions were selected for further analysis, including long-read PacBio sequencing for founder individuals and a founder haplotype prediction test. A similarity analysis at a genomic level and for loci encoding transcription factors predicted to interact with selected sequences inside the associated loci were accomplished. A total of 1,181 polymorphisms were associated with URA. All associated polymorphisms were located on chromosome 14 between 32.9 and 36.6 Mb. The most significant polymorphism was chr14:36,411,266, a G/T transversion. The same associated region was identified in population three. Polygenic epistasis was determined as not predominant for the presentation of URA. Based on the haplotype mosaic probability estimation, cases display a higher probability of inheriting the ACI allele. The long-read sequencing analysis showed the presence of an Erv insertion inside the intron one of the *KIT* gene located inside the associated region. The Erv insertion comprises one Erv sequence and two Ltr sequences located downstream and upstream of the former. No Erv insertion was identified for the founder strain BN. For ACI and HSRA, only one Ltr sequence was identified. One hundred and seven genes encoding TFs that recognize binding sites on the Erv insertion were analyzed for sequence similarity against the reference HSRA. The TF similarity score analysis for the interaction genotype and phenotype showed significance after FDR correction for 20 TFs, including AHR, HNF1B, JUNB, RARG, and RXRA. A mechanism identifying URA as a threshold phenotype is suggested in HS rats. It implies the existence of a minimum threshold for the final number of nephrons and kidney associated structures required for stalling the apoptotic process of the metanephric rudiments. Animals exhibiting a quantitative cumulative defect would express URA, being this malformation identified as a phenotype with decreased penetrance in the assessed population of HS rats. All these processes are described as mediated by KIT and TFs able to interact with sequences of the Erv insertion.

## 2. Introduction

Human unilateral renal agenesis (URA) is a congenital urinary tract malformation, and affected individuals have only one kidney (Ara et al., 2020). Human URA occurs at around 1 in 500-1,000 births. It is often an asymptomatic developmental defect. Nevertheless, URA is associated with additional genitourinary anomalies (Westland et al., 2013; Ahmed et al., 2017; Nikam et al., 2018) and a higher risk for hypertension, proteinuria, and progression to chronic kidney disease (González et al., 2005; Westland et al., 2014). URA has a substantial genetic component, with offspring and relatives of affected individuals having a significantly increased risk for URA (McPherson, 2007; Solberg Woods et al., 2010).

URA is present in a variety of species. Among rats, the inbred ACI strain shows a high rate of URA compared with other inbred strains such as BN (Shull et al., 2006). Shull et al. (2006) and Becker et al. (2015) reported that both male and female rats from the ACI strain show a URA incidence between 5 and 15%. The associated locus for this phenotype was mapped to a region on chromosome 14 (RNO14). This locus is called Renag1 (Renal agenesis 1), the major contributing factor associated with URA in ACI crosses. Shull et al. (2006) identified the Renag1 locus as a 14.4-Mb interval, delimited by D14Rat50 and D14Rat12, two microsatellites. Shull et al. (2006) also reported that ACI alleles in this locus cause URA by acting as an incompletely dominant and incompletely penetrant factor. Becker et al. (2015) performed fine mapping of Renag1 and identified a 379 kilobase (kb) interval. This locus contained only one protein-coding gene, the *KIT* (*KIT Proto-Oncogene, Receptor Tyrosine Kinase*) gene. Becker et al. (2015) identified *KIT* and *KIT* ligand expression in the nephric duct, a crucial tissue for kidney organogenesis. An endogenous retrovirus-derived long terminal repeat within the first intron of *KIT* could be the causative polymorphism for URA (Erv insertion). More recently, Ara et al. (2020) used a CRISPR/Cas9 system to remove one of the Ltr sequences of the Erv insertion and reported suppression of URA phenotype in ACI rats.

Heterogeneous stock (HS) rats constitute an outbred population created in 1984 by interbreeding eight inbred strains, including ACI (Solberg Woods and Palmer, 2019). As part of a large multi-center study of drug abuse-related behavioral traits (www.ratgenes.org). Showmaker et al. (2020) used a model of congenital abnormalities of the kidney and urogenital tract or CAKUT named HSRA. This inbred strain was generated by phenotypic selection from a high URA incidence HS family from the same population used in the present analysis. The latter ensures the presence of shared QTLs and penetrance modifiers for URA between HSRA and the assessed HS population. Applying inbreeding and phenotypic-driven selection, HSRA offspring developed a URA susceptibility between 50 and 75%. This increased susceptibility to URA must result from an increased frequency of deleterious alleles involved in the presentation of URA.

The present analysis aimed to identify genomic regions associated with URA controlling for case-control imbalance in an HS rat population. URA was recorded in more than 5,000 HS rats and subsequently genotyped at three million and a half polymorphisms across the genome. This information allows to perform a genome-wide association (GWA) analysis for URA and to conduct additional genetic analysis to identify loci able to explain incomplete penetrance and observed differences in the incidence of URA across strains and populations. Two other rat populations were evaluated to identify and validate any associated loci. A “polygenic epistasis” test applying a gene by ancestry association assessment was performed. The associated locus was additionally evaluated using long-read PacBio sequencing, including animals from each of the original HS founder strains and HSRA individuals. This procedure aims at identifying recognizable genomic elements present in this region, such as the previously identified Erv insertion. Through the identification of genomic elements present in this region, a kidney-related transcription factor (TF) prediction for the Erv insertion was performed in order to identify additional loci showing evidence of genomic selection in HSRA rats, potentially able to explain its increased URA incidence.

## 3. Materials and methods

Research protocols were approved by the Institutional Animal Care and Use Committee Protocol of the Universities of Michigan (PRO00008758) and at Buffalo (PSY03092Y), and the Tennessee Health Science Center (20-0131). The HS colony was established in 1984 by interbreeding eight inbred founder strains: ACI/N, BN/N, BUF/N, F344/N, M520/N, MR/N, WKY/N, and WN/N (Gileta et al., 2020). The rats used in this study were from generations 73 to 80. Breeders were fed Teklad 5010 diet *ad libitum* (Envigo, Madison, Wisconsin). All rats were part of a multi-site project examining multiple behavioral phenotypes related to drug abuse (https://ratgenes.org/). A total of 5,585 male and female HS rats were analyzed; these rats were assessed at three different phenotyping centers in the following manner: 1,522 at the University of Michigan, 1,596 at the University at Buffalo, and the remaining 2,467 at the University of Tennessee Health Sciences Center. The mean age of the euthanized rats was 89.93, 195.40, and 105.08 days old, respectively.

### 3.1 Genotyping

A total of 3,513,321 markers were genotyped using genotyping-by-sequencing (Parker et al., 2016; Gileta et al., 2020), excluding markers on sexual chromosomes. The Rnor_6.0 assembly was used as the reference genome. A pruned genotype dataset was generated using PLINK (Purcell, 2020). This dataset included 134,918 polymorphisms across the genome, and it was generated by using windows of 50 kb and a threshold r2 equal to 0.98. Genotypic data is available on UC San Diego Library Digital Collections https://doi.org/10.6075/J0028RR4

### 3.2 Phenotyping

Animals were euthanized by using pentobarbital overdose and posterior decapitation. Chitre et al. (2020) present additional details about breeding and housing. Animals were assessed for renal agenesis.

### 3.3 Association analysis

A logistic regression analysis was performed using the R package aod (Lesnoff and Lancelot, 2012) to identify significant covariables for the variable URA, performing a comparison against the null model. Age and factors such as sex, center, and batch were recorded and assessed as covariables. The final association model included only the polymorphism being tested as a covariable in the association model. The R package SAIGEgds (Zhou et al., 2018) was used for the association analysis. This package performs a scalable and accurate generalized mixed model association test. It applies the saddlepoint approximation to calibrate the distribution of score test statistics accounting for case-control imbalance. Permutation analysis was performed to determine the association threshold for this analysis by performing 1,000 permutations. The adjusted p-value threshold for the association analysis was equal to 5.6 (-log10). The R package “CMplot” v3.3.0 (LiLin-Yin, 2017) was used to graph p-value distributions.

### 3.4 Additional association analysis

Two additional datasets were used as validation tests. Population two included 1,577 rats genotyped for 7,425,889 markers. The case-control imbalance was equal to 1:174 (9 cases). The population three included 1,407 rats. This population was genotyped for 254,932 markers, and the case-control ratio was equal to 1:38 (37 cases). The accession number for this dataset is E-MTAB-2332 from ArrayExpress (www.ebi.ac.uk). A permutation analysis to determine the association threshold for these populations, including 1,000 permutations, was determined. The association threshold for population two and population three was 5.1 and 5.2, respectively.

### 3.5 Gene by ancestry association

The Python package GxTheta was used to perform a gene by ancestry association (Van Rossum and Drake Jr, 1995; Rau et al., 2020) in a pruned genotypic dataset for each parental strain separately. The pruned dataset was used. The association threshold was calculated by using permutation and then this threshold was divided by the number of the tested founder strains (eight). The final computed significance threshold was equal to 6.5 (-log10). The R package “CMplot” v3.3.0 (LiLin-Yin, 2017) was used to graph the association results.

### 3.6 Founder haplotype mosaic and exclusive founder alleles

The founder haplotype mosaic was reconstructed using the unpruned dataset. The haplotype estimation was performed for controls and cases independently. Additionally, founder haplotype mosaic probabilities for the most highly associated segment between chr14:32-37 Mb were estimated. The R package QTL2 was used for this estimation (Karl et al., 2019), and the results were graphed using the R package CMplot (LiLin-Yin, 2017) and Ggplot2 (Wickham, 2016). Exclusive founder alleles were identified for the associated locus.

### 3.7 Long read PacBio sequencing

Guidelines from the “extracting HMW DNA from animal tissue using TissueRuptor” and “Preparing whole genome and metagenome libraries using SMRTbell®” protocols were used (www.pacbio.com). Samples of ∼25 mg of tissue from liver, small intestine, tail, or spleen tissue from all eight founder strains and the HSRA strain were processed with the Pacific Biosciences (PacBio) Nanobind tissue kit. DNA libraries were prepared with PacBio SMRTbell prep kit 3.0. Extracted DNA was sequenced with PacBio Sequel IIe Circular Consensus Sequencing (CCS) equipment to obtain 15x mean coverage high fidelity (HiFi) reads (Wenger et al., 2019). The Institute for Genomic Medicine IGM at the University of California San Diego performed DNA library construction and sequencing using a PacBio Sequel IIe system. Samtools was used to aligning reads to the Rnor_6.0 assembly (Li et al., 2009; Li, 2018). Reads mapping to the associated locus were visualized using the IGV software (Robinson et al., 2011, 2017, 2023; Thorvaldsdóttir et al., 2012).

### 3.8 Transcription factor binding site prediction

The software PROMO was used to predict putative transcription factor binding sites (TFBS) for the Erv insertion sequences (Messeguer et al., 2002); TFs for animals available in the TRANSFAC database version 8.3 were included in the analysis. The maximum matrix dissimilarity rate for predicting TFBSs on the target sequences was 10% (90% similarity). This list of TFs was used as a query on the Mouse Genome Informatics website (Bult CJ, Blake JA, Smith CL, Kadin JA, Richardson JE, 2019), and TFs with expression in tissues and structures associated with the kidney were retained. A total of 89 kidney-associated tissues and structures (Table 1) were selected for this analysis. The final list of TFs expressed and associated with kidney tissues and structures was included in a similarity analysis.

**Table 1.**
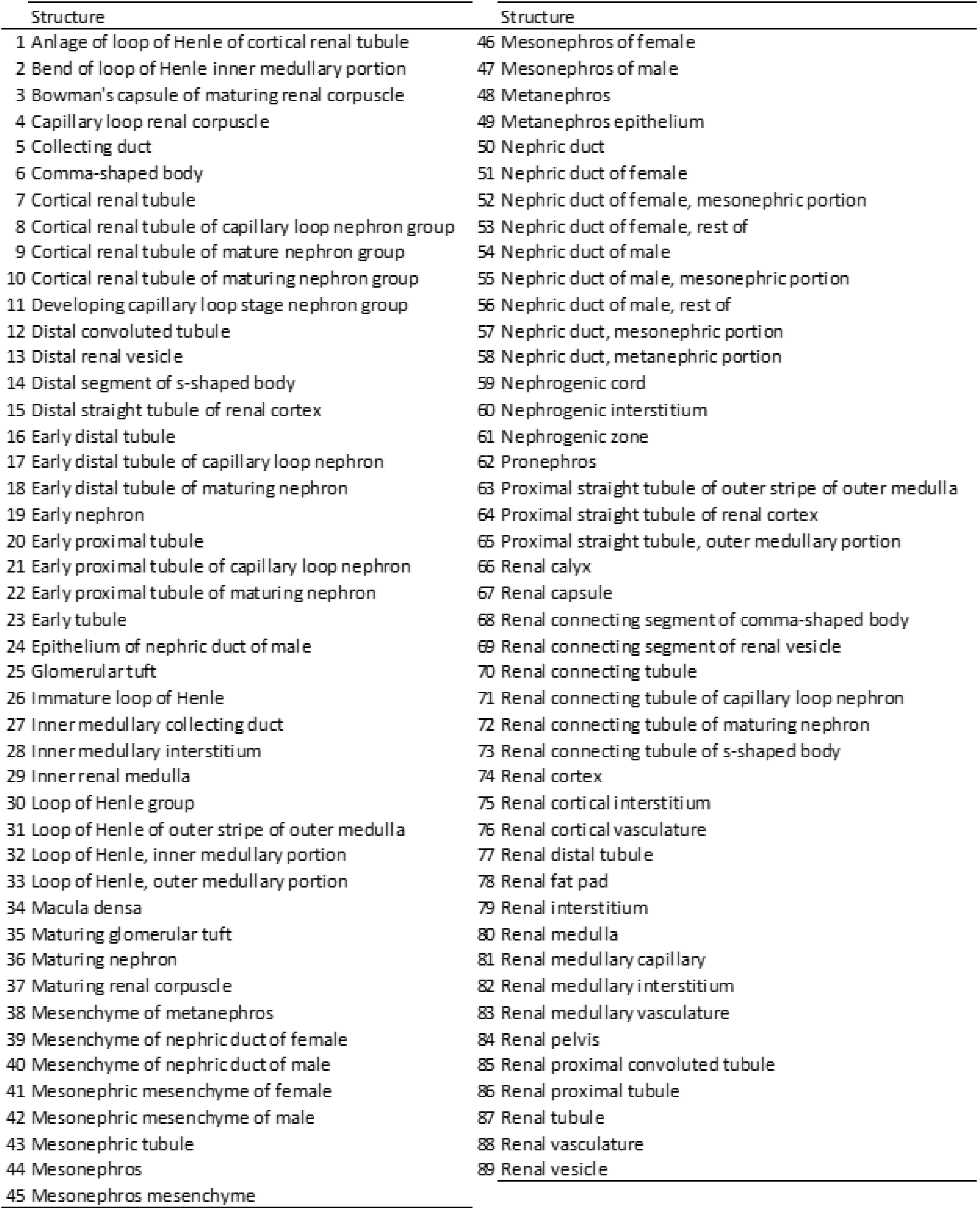
Selected tissues and structures associated with the kidney used for screening of transcription factors.

| Structure | Structure |
| --- | --- |
| 1 Anlage of loop of Henle of cortical renal tubule | 46 Mesonephros of female |
| 2 Bend of loop of Henle inner medullary portion | 47 Mesonephros of male |
| 3 Bowman's capsule of maturing renal corpuscle | 48 Metanephros |
| 4 Capillary loop renal corpuscle | 49 Metanephros epithelium |
| 5 Collecting duct | 50 Nephric duct |
| 6 Comma-shaped body | 51 Nephric duct of female |
| 7 Cortical renal tubule | 52 Nephric duct of female, mesonephric portion |
| 8 Cortical renal tubule of capillary loop nephron group | 53 Nephric duct of female, rest of |
| 9 Cortical renal tubule of mature nephron group | 54 Nephric duct of male |
| 10 Cortical renal tubule of maturing nephron group | 55 Nephric duct of male, mesonephric portion |
| 11 Developing capillary loop stage nephron group | 56 Nephric duct of male, rest of |
| 12 Distal convoluted tubule | 57 Nephric duct, mesonephric portion |
| 13 Distal renal vesicle | 58 Nephric duct, metanephric portion |
| 14 Distal segment of s-shaped body | 59 Nephrogenic cord |
| 15 Distal straight tubule of renal cortex | 60 Nephrogenic interstitium |
| 16 Early distal tubule | 61 Nephrogenic zone |
| 17 Early distal tubule of capillary loop nephron | 62 Pronephros |
| 18 Early distal tubule of maturing nephron | 63 Proximal straight tubule of outer stripe of outer medulla |
| 19 Early nephron | 64 Proximal straight tubule of renal cortex |
| 20 Early proximal tubule | 65 Proximal straight tubule, outer medullary portion |
| 21 Early proximal tubule of capillary loop nephron | 66 Renal calyx |
| 22 Early proximal tubule of maturing nephron | 67 Renal capsule |
| 23 Early tubule | 68 Renal connecting segment of comma-shaped body |
| 24 Epithelium of nephric duct of male | 69 Renal connecting segment of renal vesicle |
| 25 Glomerular tuft | 70 Renal connecting tubule |
| 26 Immature loop of Henle | 71 Renal connecting tubule of capillary loop nephron |
| 27 Inner medullary collecting duct | 72 Renal connecting tubule of maturing nephron |
| 28 Inner medullary interstitium | 73 Renal connecting tubule of s-shaped body |
| 29 Inner renal medulla | 74 Renal cortex |
| 30 Loop of Henle group | 75 Renal cortical interstitium |
| 31 Loop of Henle of outer stripe of outer medulla | 76 Renal cortical vasculature |
| 32 Loop of Henle, inner medullary portion | 77 Renal distal tubule |
| 33 Loop of Henle, outer medullary portion | 78 Renal fat pad |
| 34 Macula densa | 79 Renal interstitium |
| 35 Maturing glomerular tuft | 80 Renal medulla |
| 36 Maturing nephron | 81 Renal medullary capillary |
| 37 Maturing renal corpuscle | 82 Renal medullary interstitium |
| 38 Mesenchyme of metanephros | 83 Renal medullary vasculature |
| 39 Mesenchyme of nephric duct of female | 84 Renal pelvis |
| 40 Mesenchyme of nephric duct of male | 85 Renal proximal convoluted tubule |
| 41 Mesonephric mesenchyme of female | 86 Renal proximal tubule |
| 42 Mesonephric mesenchyme of male | 87 Renal tubule |
| 43 Mesonephric tubule | 88 Renal vasculature |
| 44 Mesonephros | 89 Renal vesicle |
| 45 Mesonephros mesenchyme |  |

### 3.9 Similarity analysis

A sequence similarity analysis was performed, including the HSRA strain as the reference sequence, since it was inbred for URA and was selected from the same HS population (Showmaker et al., 2020). HSRA is a valuable reference to look for sequence similarity to uncover potential penetrance modifiers involved in increasing susceptibility to URA. Out of 5,585 individuals, 5,576 animals were selected for the similarity analysis. An in-house Java script was used to determine sequence similarity by applying the Jaccard Index (Besta et al., 2020), comparing all individuals against the HSRA strain. The pruned dataset was used for this analysis. The Jaccard similarity score is expressed as a percentage and computed for each pair of data samples in the following manner:

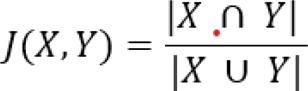

Where X and Y are sample sets (Besta et al., 2020). The Jaccard similarity score represents the percentage of shared elements between both sets, HS versus HSRA rats. Similarity score estimation was performed in the following way:

1. Whole genome excluding chromosome 14.
2. Analysis for chromosome 14.
3. Assessment for the associated locus.
4. Evaluation for loci harboring selected TFs.

The genomic locations for the selected TF genes were obtained using Biomart from Ensembl (Zerbino et al., 2018). Table 2 presents the 107 TF gene genomic locations. Gene location was extended by 100 kb upstream and downstream to include and analyze potential regulatory sequences (Gaffney et al., 2012). Genes located on chromosome X were excluded from the analysis. For the genic similarity score, grouping was performed by genotype at the most significantly associated marker for URA (chr14:36,411,266) classified by phenotype (control versus cases). Genic similarity scores for all individuals were estimated. ANOVA was used to identify significant TFs for the interaction between phenotype and genotype, and for the interaction phenotype*genotype. Test p-values for this analysis were corrected using a false discovery rate (FDR) equivalent to an alpha of 10% using the Benjamini Hochberg appraoch. Only the interaction genotype*phenotype was assessed because an analysis by genotype or phenotype independently would not allow to identify loci able to explain differences between cases and controls given the genotype at the associated URA locus. This analysis aims at identifying additional loci able to explain incomplete penetrance in this population, identifying genes encoding TFs with statistical differences for means when assessing the interaction genotype*phenotype.

**Table 2.**
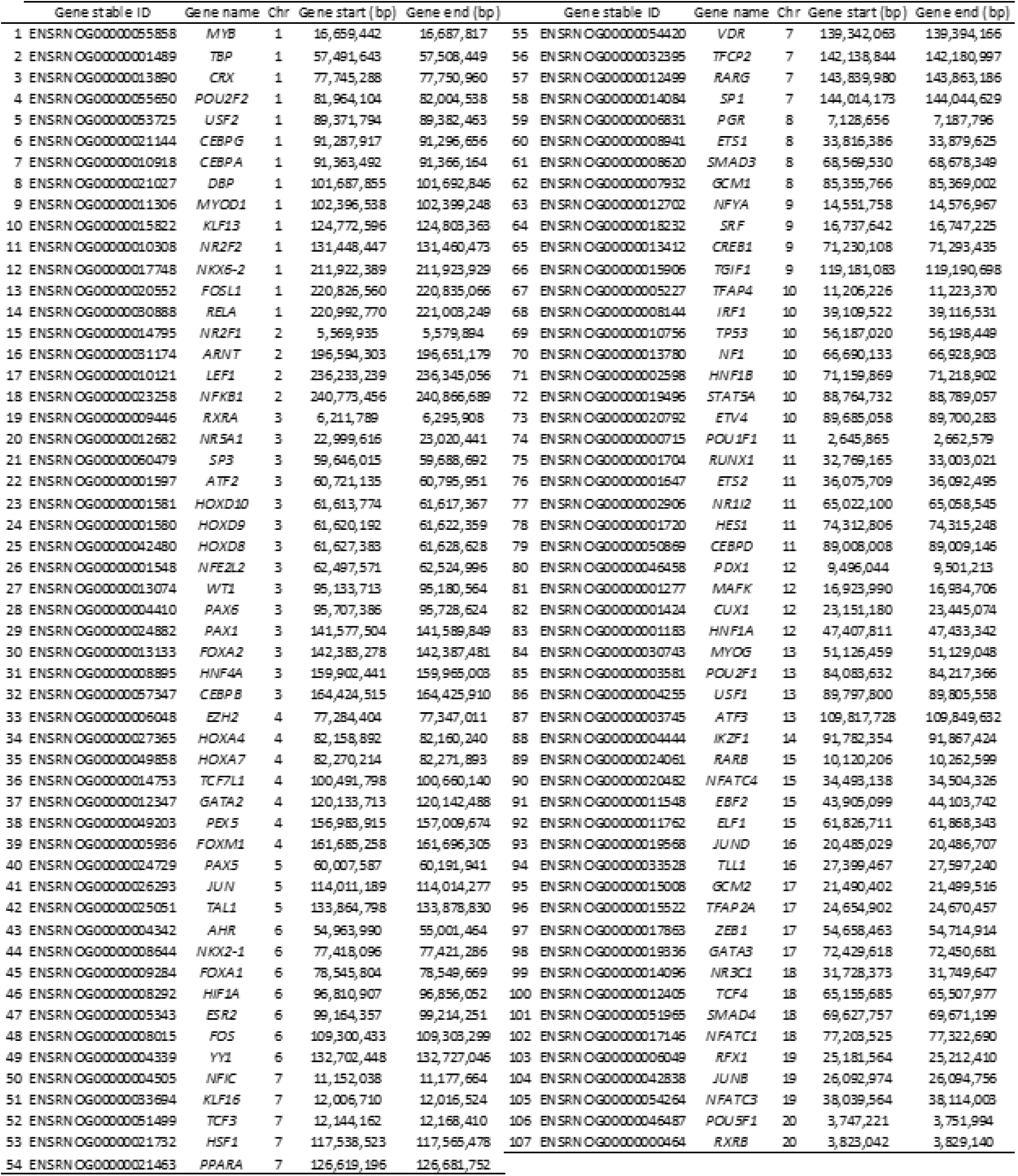
Selected genes and their genomic location. The target region used for the similarity analysis was obtained by including 100 kbs up and downstream of the shown gene location. The Rnor_6.0 assembly was used as the reference genome.

## 4. Results

### 4.1 Association analysis

The association analysis was performed including a total of 110 cases (case-control ratio equal to 1:50.7). Supplementary Table 1 and Figure 1 show the association analysis results. A total of 1,181 polymorphisms were associated with URA. All these polymorphisms were on chromosome 14 between 32.9 and 36.6 Mb. The most significant polymorphism was chr14:36,411,266, a G/T transversion. The proportion of cases by genotype was 0.48% (15 out of 3,127), 3.5% (75 out of 2,060), and 6.49% (20 out of 288) for G/G, G/T, and T/T, respectively.

**Figure 1a.**
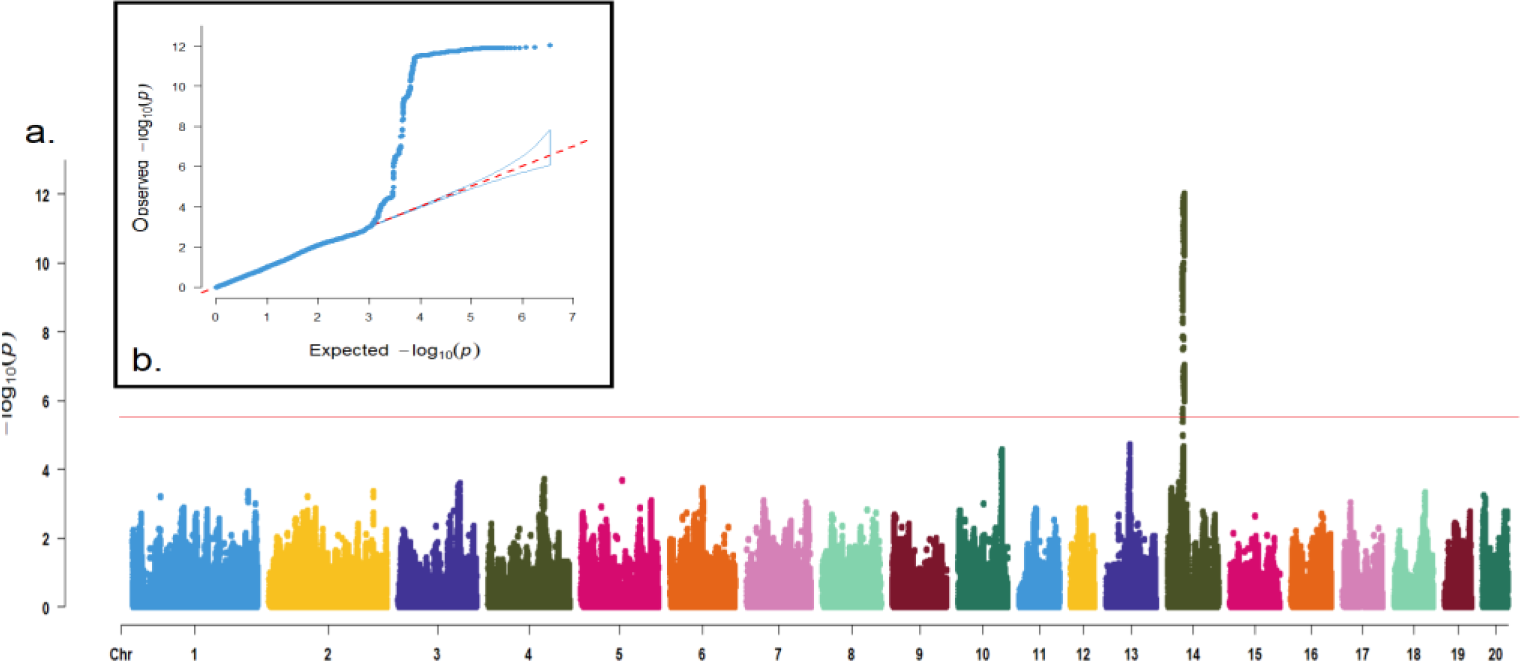
Whole-genome association analysis results for URA. A total of 5,585 individuals were genotyped for 3,513,321 markers. The analysis was performed using the R package SAIGEgds. The red line represents the corrected p-value threshold obtained using permutation (-log10 equal to 5.6); 1b. Qqplot including the distribution of observed and expected p-values for the association analysis.

A detail of the most highly significantly associated region on chromosome 14, between positions 32.9 and 36.6 Mb, is presented in Figure 2. Fifty-three genes and one pseudogene are inside this region. Twelve lincRNAs, 36 protein-coding genes, two snoRNAs, one miRNA, one rRNA (*5S_RRNA*), and one snRNA (*U6*) were identified. Between the protein-coding genes, the following were identified: *Insulin Like Growth Factor Binding Protein 7* (*IGFBP7*), *KIT* (shown in red), *Transmembrane Protein 165* (*TMEM165*), and *Ubiquitin Specific Peptidase 46* (*USP46*).

**Figure 2.**
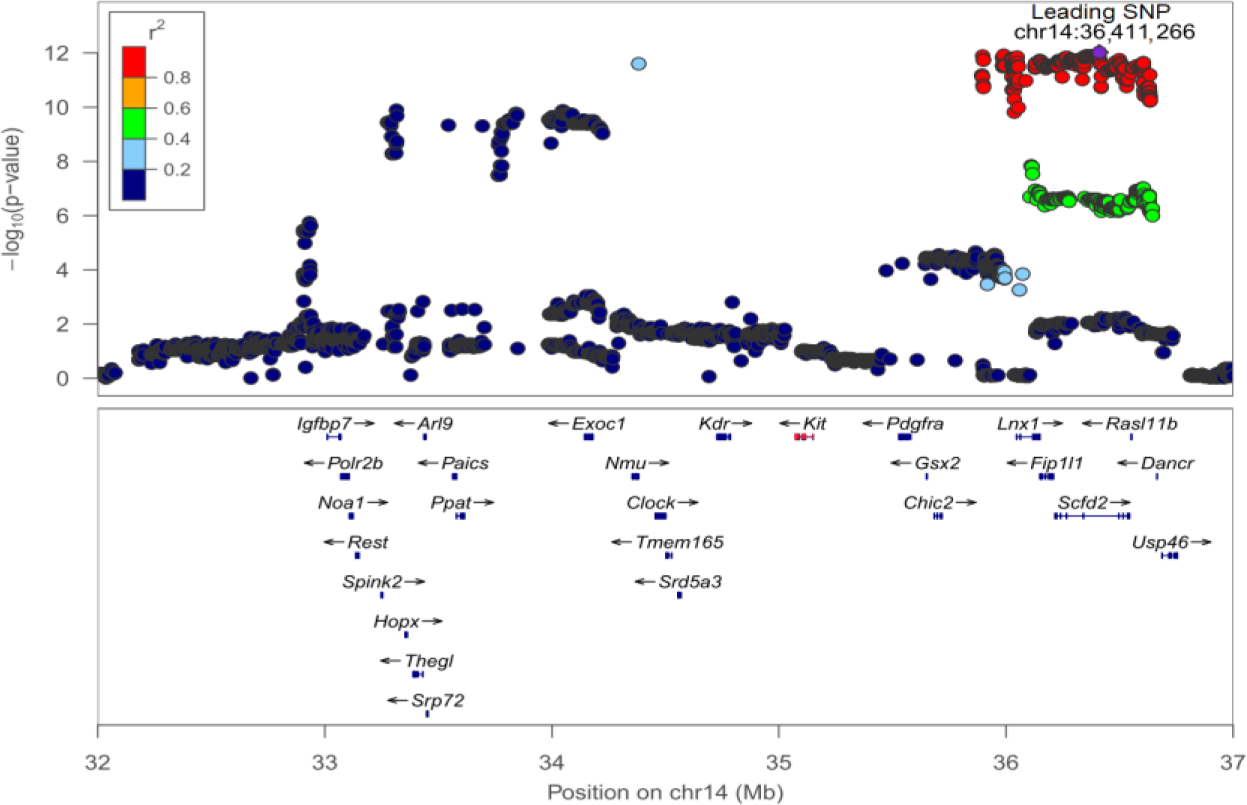
Regional association plot for the most significantly associated locus located on chr14:32.9-36.6 Mb; The analysis was performed using the R package SAIGEgds; Dot color represents the amount of linkage disequilibrium (r2) between each SNP and the top associated SNP (chr14:36,411,266). Gene distribution on chr14:32.9-36.6 Mb is presented. The *KIT* gene is shown in red.

Since the number of associated markers in the same region was high, another association analysis was performed, including the most significantly associated polymorphism, chr14:36,411,266, as a covariable in the model. The polymorphism chr14:36,411,266 was able to account for all the variability present on chr14:32.9-36.6 since no marker remained as significantly associated with URA.

### 4.2 Additional association analysis

Only one additional dataset, population 3, showed replication of the present results (Figure 3, Table 3). The segment Chr14:33,388,086-34,490,097 Mb harbored 30 significant polymorphisms.

**Figure 3.**
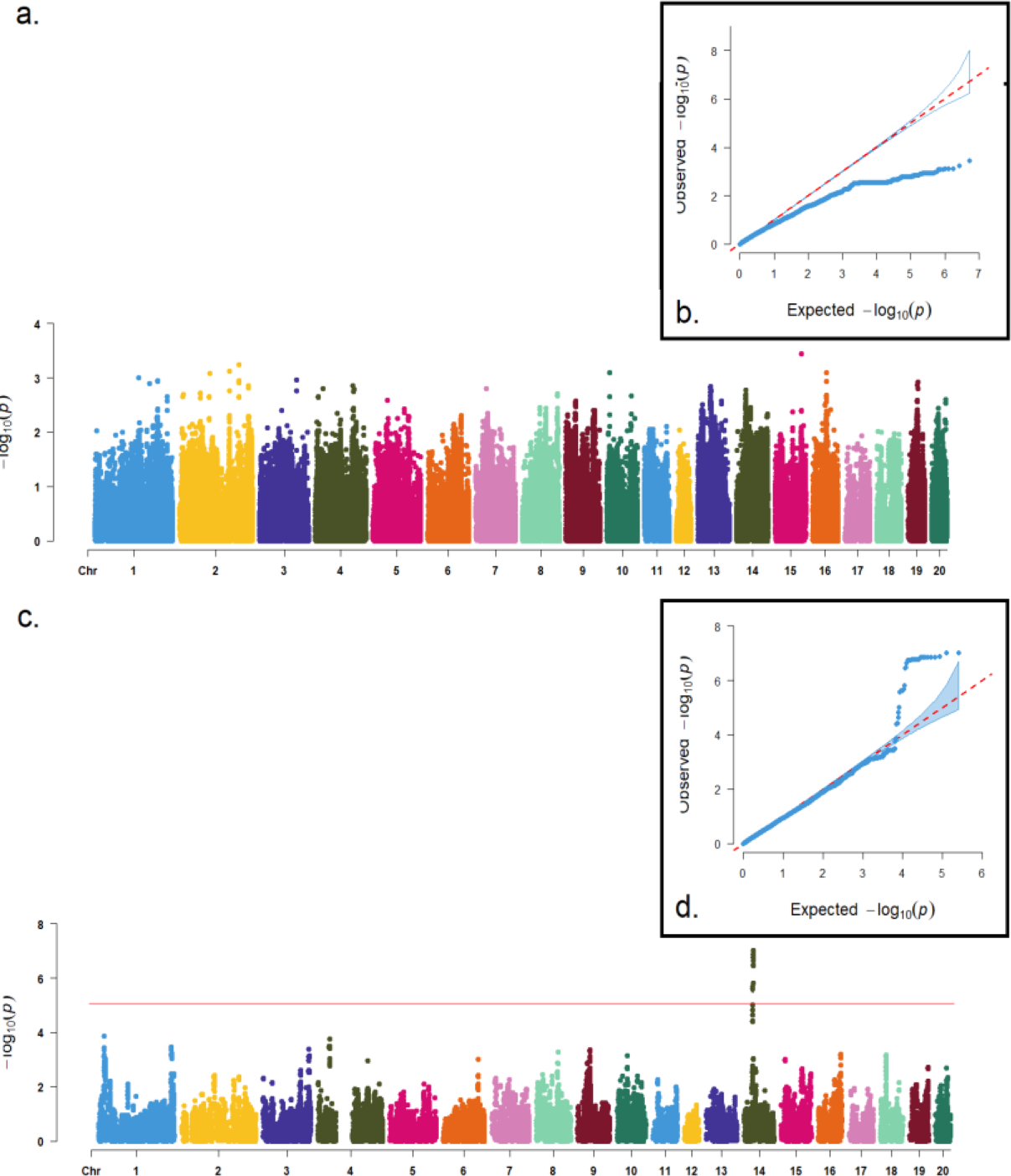
Additional association analyses for URA. 3a. Association results for the population two. This population included 1,577 rats genotyped for 7,425,889 markers (case:control ratio = 1:174). 3b. Qqplot includes the distribution of observed and expected p-values for the association analysis performed on the population two. 3c. Association results for the population three. It included 1,407 rats, genotyped for 254,932 markers (case:control ratio = 1:38); 3d. Qqplot including the distribution of observed and expected p-values for the association analysis performed on this population.

**Table 3.** Additional association analyses for URA. Results for population three; 1,407 rats were included and genotyped for 254,932 markers. The case-control ratio was equal to 1:38.

| Marker | Chromosome | Position | Log | Marker | Chromosome | Position | Log |
| --- | --- | --- | --- | --- | --- | --- | --- |
| chr14:33388086 | 14 | 33,388,086 | 7.0 | chr14:34042826 | 14 | 34,042,826 | 6.8 |
| chr14:33434566 | 14 | 33,434,566 | 7.0 | chr14:34046305 | 14 | 34,046,305 | 6.8 |
| chr14:33456622 | 14 | 33,456,622 | 6.9 | chr14:34053556 | 14 | 34,053,556 | 6.8 |
| chr14:33457630 | 14 | 33,457,630 | 6.9 | chr14:34054202 | 14 | 34,054,202 | 6.7 |
| chr14:33463100 | 14 | 33,463,100 | 6.9 | chr14:34151694 | 14 | 34,151,694 | 6.6 |
| chr14:33473958 | 14 | 33,473,958 | 6.9 | chr14:34156236 | 14 | 34,156,236 | 6.5 |
| chr14:33562929 | 14 | 33,562,929 | 6.9 | chr14:34156512 | 14 | 34,156,512 | 6.5 |
| chr14:33826982 | 14 | 33,826,982 | 6.9 | chr14:34168554 | 14 | 34,168,554 | 5.8 |
| chr14:33869364 | 14 | 33,869,364 | 6.9 | chr14:34178218 | 14 | 34,178,218 | 5.7 |
| chr14:33877981 | 14 | 33,877,981 | 6.8 | chr14:34178683 | 14 | 34,178,683 | 5.6 |
| chr14:33982434 | 14 | 33,982,434 | 6.8 | chr14:34179113 | 14 | 34,179,113 | 5.6 |
| chr14:33984825 | 14 | 33,984,825 | 6.8 | chr14:34179426 | 14 | 34,179,426 | 5.6 |
| chr14:33985056 | 14 | 33,985,056 | 6.8 | chr14:34182945 | 14 | 34,182,945 | 5.6 |
| chr14:34025204 | 14 | 34,025,204 | 6.8 | chr14:34183147 | 14 | 34,183,147 | 5.6 |
| chr14:34039143 | 14 | 34,039,143 | 6.8 | chr14:34490097 | 14 | 34,490,097 | 5.6 |

### 4.3 Gene by ancestry association

Table 4 and Figure 4 show the association results for GxTheta. Seven polymorphisms were above the association threshold, including markers chr10:70,904,834 (ACI), chr13:27,658,836 and chr14:108,012,542 (BN), chr13:27,658,836 and chr14:108,012,542 (BUF), chr16:4,664,758 (MR) and chr7:23,652,958 (WN).

**Figure 4.**
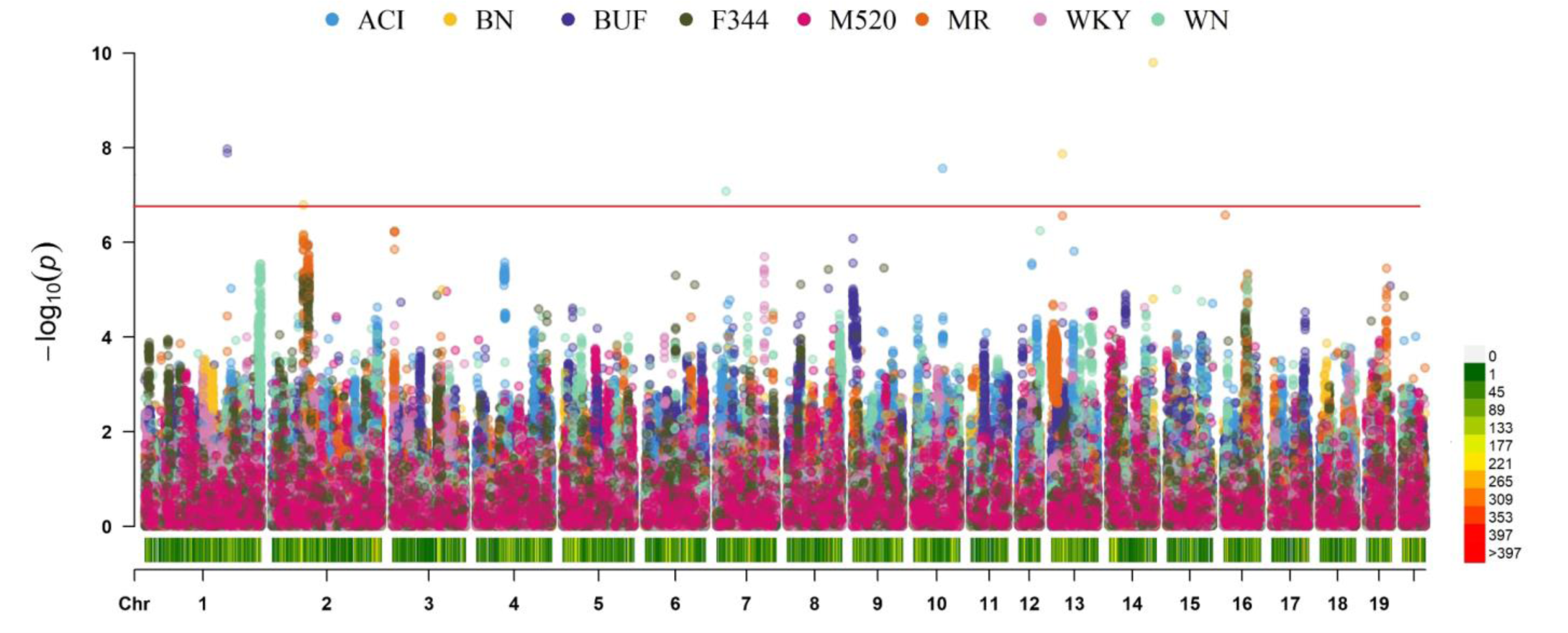
Multitrack Manhattan plot for gene by ancestry association. It shows results from the marker-founder strain interaction. Each dot was color-coded based on the tested founder strain. The solid red line represents the association threshold (-log10 equal to 6.5) calculated by dividing an association threshold acquired by permutation by the number of founder strains (eight). Polymorphism density is also presented in the bottom region of this graph.

**Table 4.**
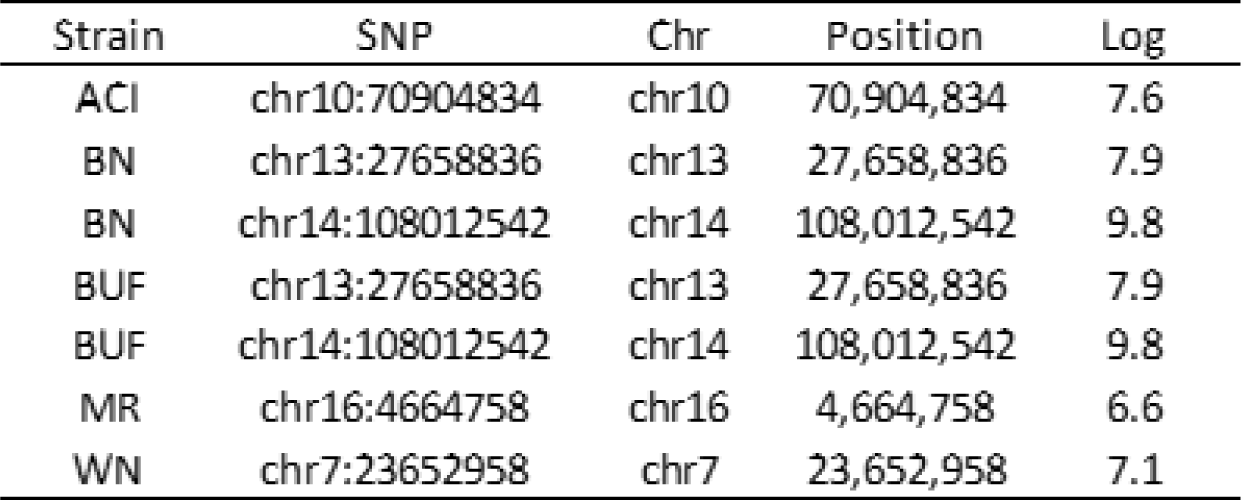
Association results from GxTheta. The association threshold was calculated by dividing the association threshold acquired by permutation by the number of founder strains (eight); the threshold was 6.5 (-log10).

### 4.4 Founder haplotype mosaic and exclusive alleles

Figure 5 and Supplementary Figure 1 show the estimation for founder haplotype composition for cases and controls. Figure 6 illustrates the association probability for exclusive alleles by founder strain and a difference in mean haplotype mosaic probability estimation for cases and controls for the associated locus 32.9-36.6 Mb. The associated locus shows an increased probability of being inherited by ACI in cases (Figure 5), the founder strain with the higher incidence of URA. Inside this region, two clusters are evident on the exclusive allele plot (Figure 6a). The most significant cluster was identified as present in ACI, BUF, and M520 simultaneously; however, this cluster was also identified as having a higher haplotypic probability of being from ACI origin when considering only cases (Figure 6b). The second cluster is an ACI-exclusive region; therefore, this region has a higher haplotypic probability of being from ACI origin for cases.

**Figure 5.**
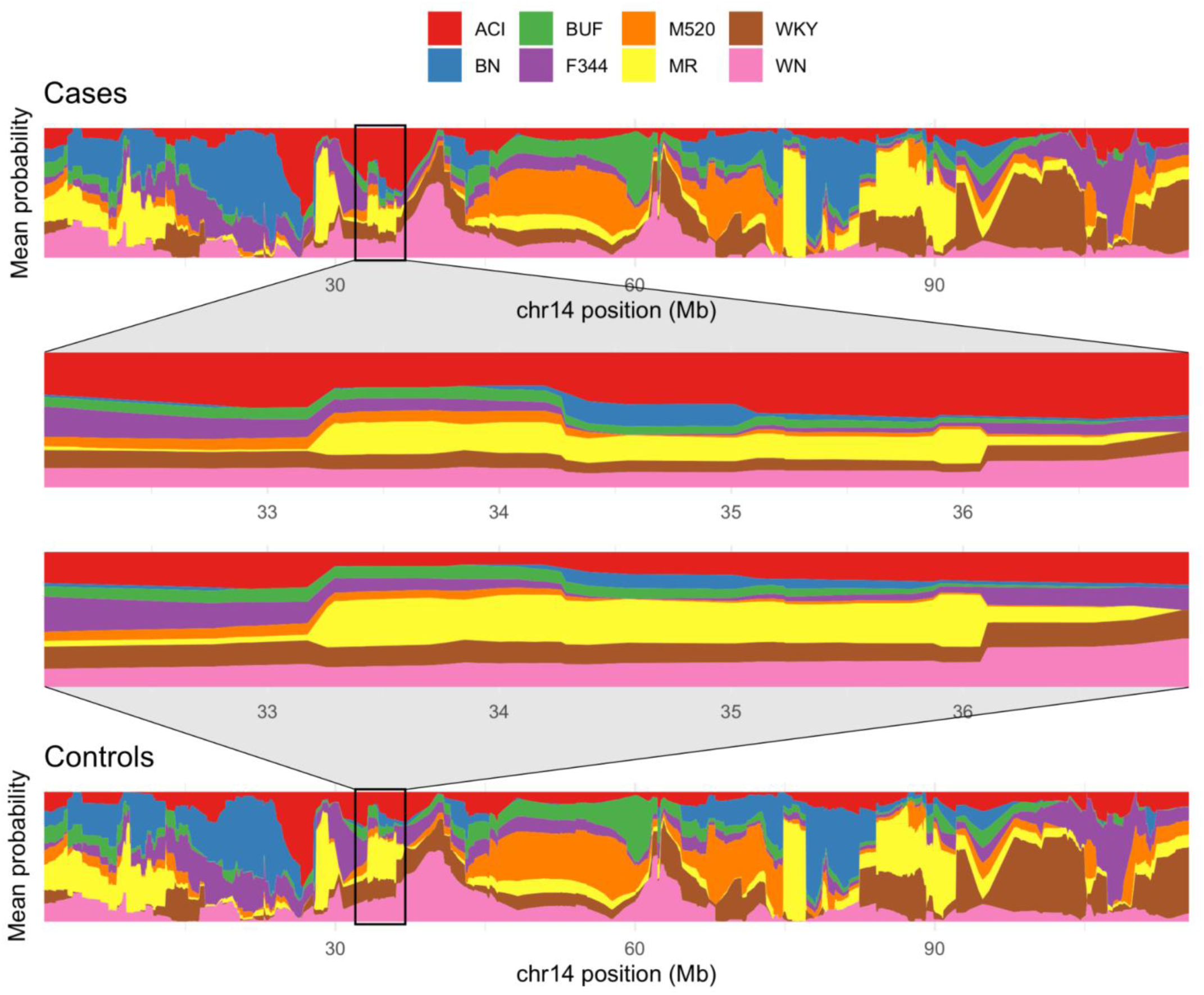
Mean probability per marker for founder haplotype mosaic for the whole chromosome 14. A comparison between cases and controls is shown. The probabilities were color-coded based on the tested founder strain. The highly associated locus located between 32.9-36.6 Mb is highlighted.

**Figure 6a.**
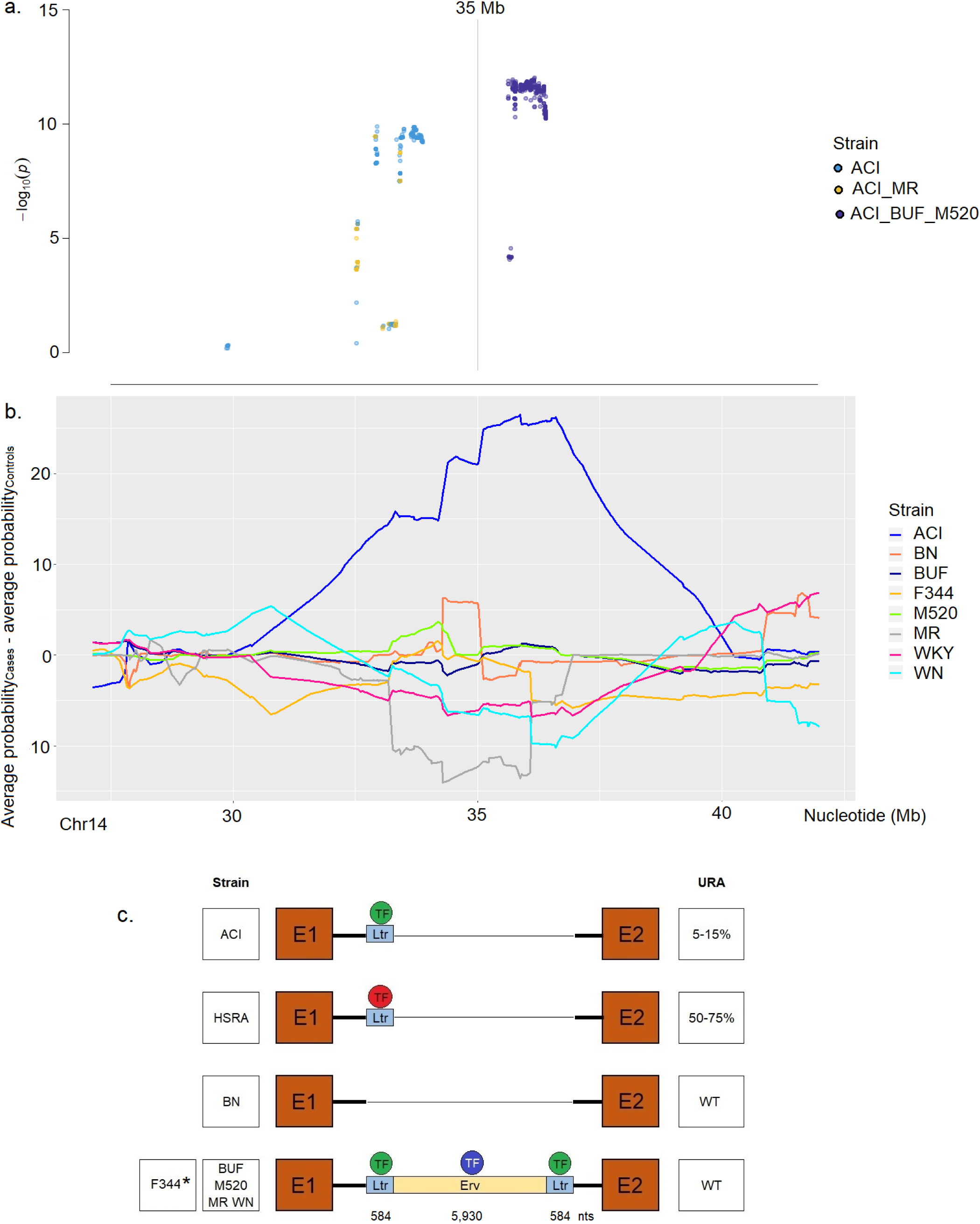
Association probability for exclusive alleles by founder strain. The locus chr14:32.9-36.6 Mb was analyzed for exclusive alleles according to founder strain origin. Dots were color-coded based on strain origin. The target locus was extended including 5 Mbs up and downstream. 6b. Mean haplotype difference for mosaic probability for cases and controls per marker. The target locus chr14:32.9-36.6 Mb is shown, including 5 Mbs up and downstream. The haplotype mosaic probability is color-coded based on founder strain origin. The location of the *KIT* gene (reported URA candidate) is chr14:35,072,131-35,149,638. 6c. Long-read sequencing results. Elements of the *KIT* intron 1 Erv insertion by founder strain are presented. *Erv element composition for F344 strain reported by Ara et al. (2020), including 7,098 nts (GenBank accession number AP012487). The theorized incomplete penetrance mechanism present in HS is described, including the Erv insertion composition by founder strain and the selected strain HSRA. The proposed mechanism for HS is shown according to reported URA incidence by strain. For ACI, the presence of only one Ltr sequence and a TF able to interact with the Ltr sequence are required to modulate URA presentation; for HSRA it was identified the existence of only one Ltr sequence and theorized the existence of an additional selected TFs (red TF). Both, Ltr and selected TF (red TF) are responsible for increasing URA penetrance; for BN, the lack of any sequence of the Erv insertion promotes the presentation of WT URA incidence; for additional founder strains, including F344, the existence of the whole Erv insertion drives WT URA incidence by counterbalancing the effect of Ltr sequence.

### 4.5 Long-read sequencing results

Figure 6c shows the long-read sequencing results. No sequences associated with the Erv insertion were identified for BN. An insertion of 584 nt corresponding to one Ltr sequence was identified inside the intron one of *KIT* for ACI and HSRA. Full Erv insertion was evident for the remaining analyzed founder strains (7,098 nts).

### 4.6 Transcription factor binding sites (TFBS) prediction

The Erv insertion is composed of one Erv sequence and two Ltr sequences located downstream and upstream of the former. A total of 107 genes encoding TFs able to recognize binding sites on the Erv insertion were analyzed, including four heteroproteins (table 5). Sixty-two TFs interact with each Ltr sequence (supplementary table 2). Some of them were CCAAT Enhancer Binding Protein Alpha (CEBPA), ETS Proto-Oncogene 1, Transcription Factor (ETS1), and GATA Binding Protein 1 (GATA1). A total of 108 TFs and four heteroproteins were identified for the Erv sequence (supplementary table 3). Some of the identified TF were: Activating Transcription Factor 2 (ATF2), Forkhead Box A1 (FOXA1), and Signal Transducer and Activator of Transcription 4 (STAT4). Both Ltrs had the exact TF prediction with slight differences in recognized sequences since they are not entirely homologous.

**Table 5.**
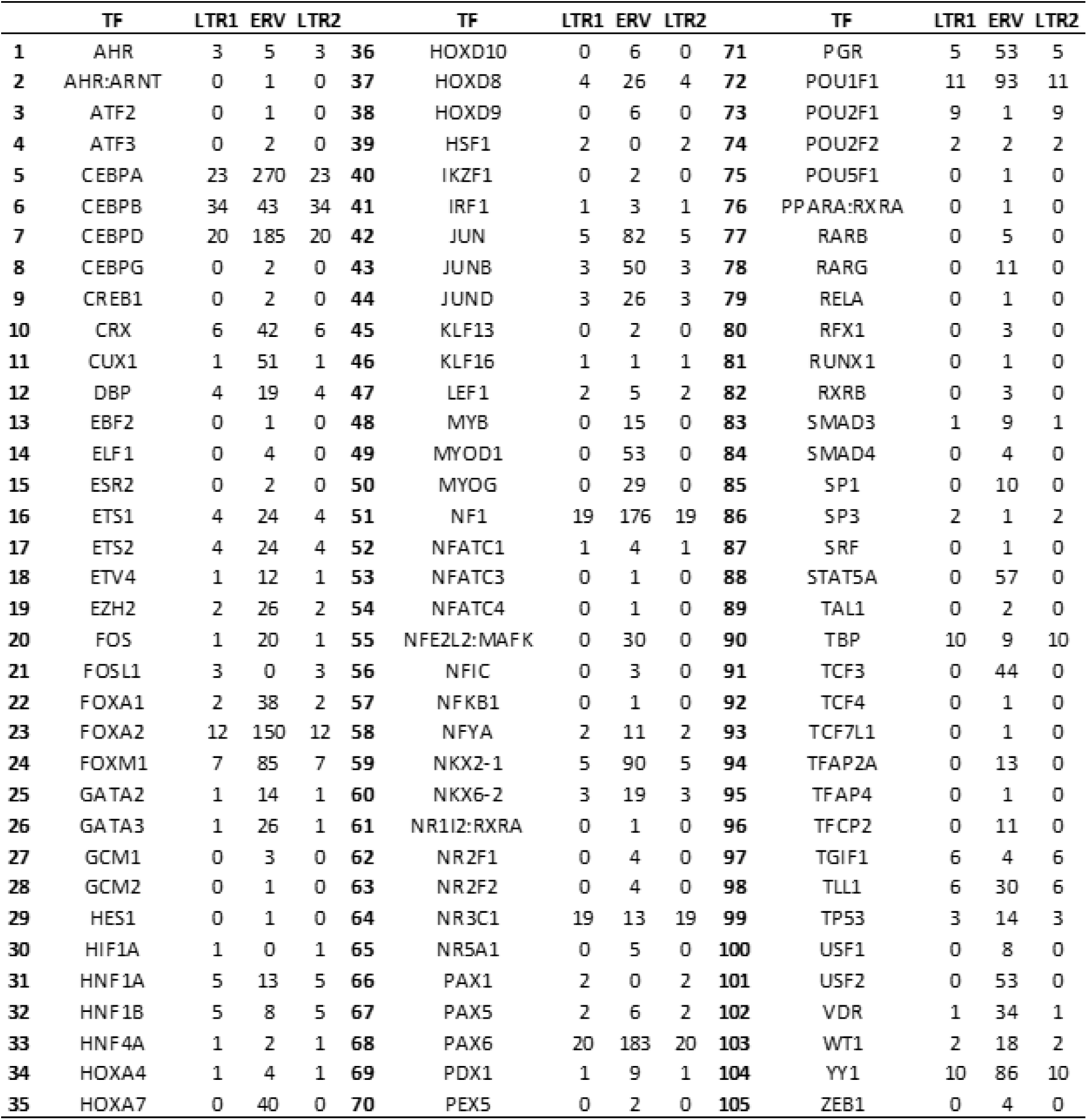
Number of binding sites identified for TFs and multiproteic TFs for the Erv insertion. The software PROMO was used to predict putative binding sites for TFs with expression in tissues and structures associated with the kidney in mice; 89 kidney-associated tissues and structures were used. TFs for animals available in the TRANSFAC database version 8.3 were included in the analysis applying a maximum matrix dissimilarity rate of 10% similarity.

### 4.7 Similarity analysis

The genomic similarity score for the interaction between genotype and phenotype is shown in Figure 7. No significant differences in the genomic similarity score analysis was found. The TF similarity score analysis for the interaction between genotype and phenotype showed significance after FDR correction for a total of 20 TFs, including Aryl Hydrocarbon Receptor (AHR), HNF1 Homeobox B (HNF1B), JunB Proto-Oncogene, AP-1 Transcription Factor Subunit (JUNB), Retinoic Acid Receptor Gamma (RARG), and Retinoid X Receptor Alpha (RXRA) (Figure 8). The whole chr14 and the associated locus chr14:32.9-36.6 Mb were identified as highly significant, showing the latter additional selection pressure in HSRA. No apparent differences between the similarity score distribution for chr14:32.9-36.6 Mb and the whole chr14 were evident, meaning that differences in chr14 seem to be driven by the associated locus.

**Figure 7.**
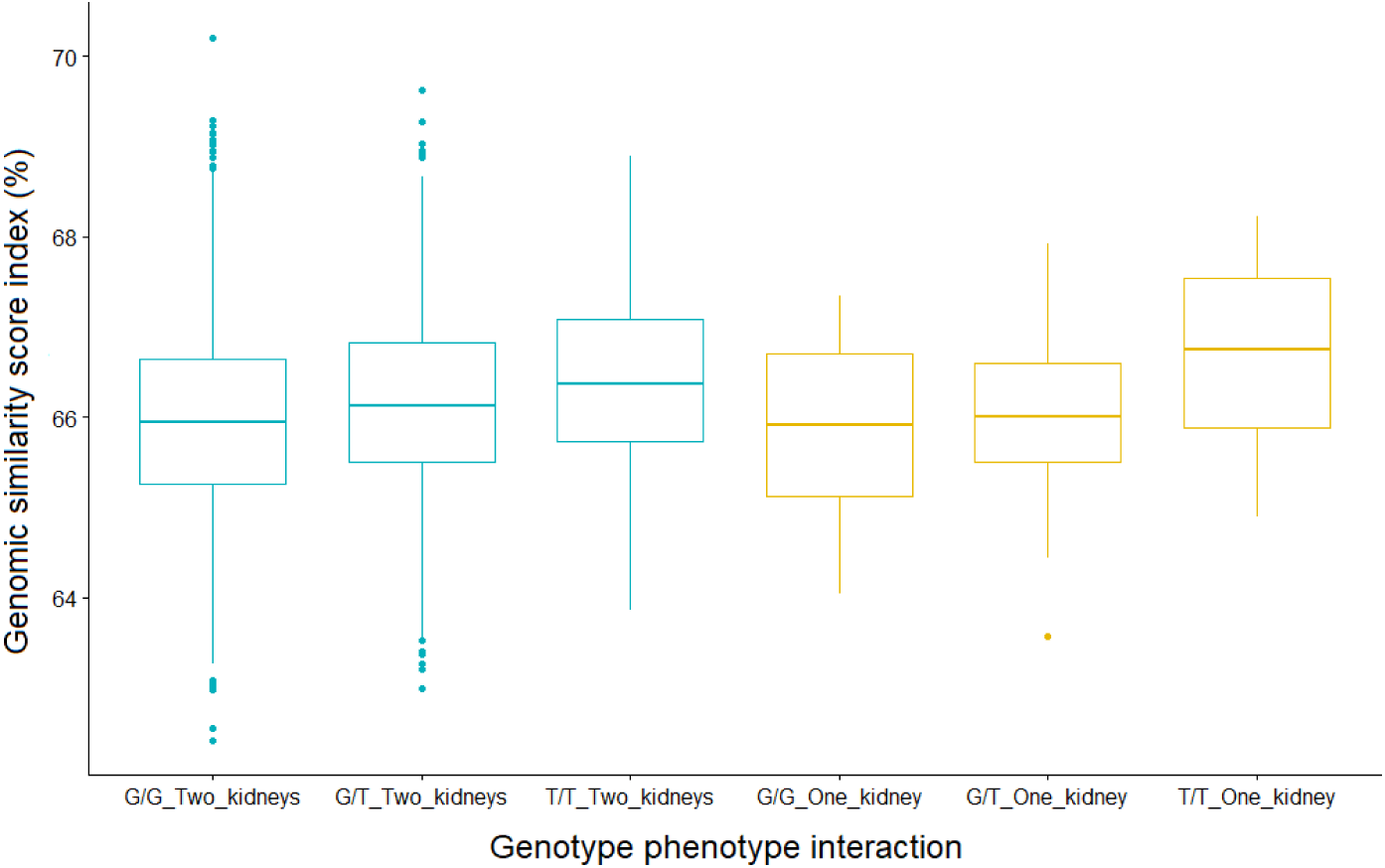
The genomic similarity score for the interaction between genotype and phenotype. Genotype grouping was performed based on the most significantly associated SNP, chr14:36,411,266. The Jaccard index was used to construct a similarity index, including the HSRA strain as the reference.

**Figure 8.**
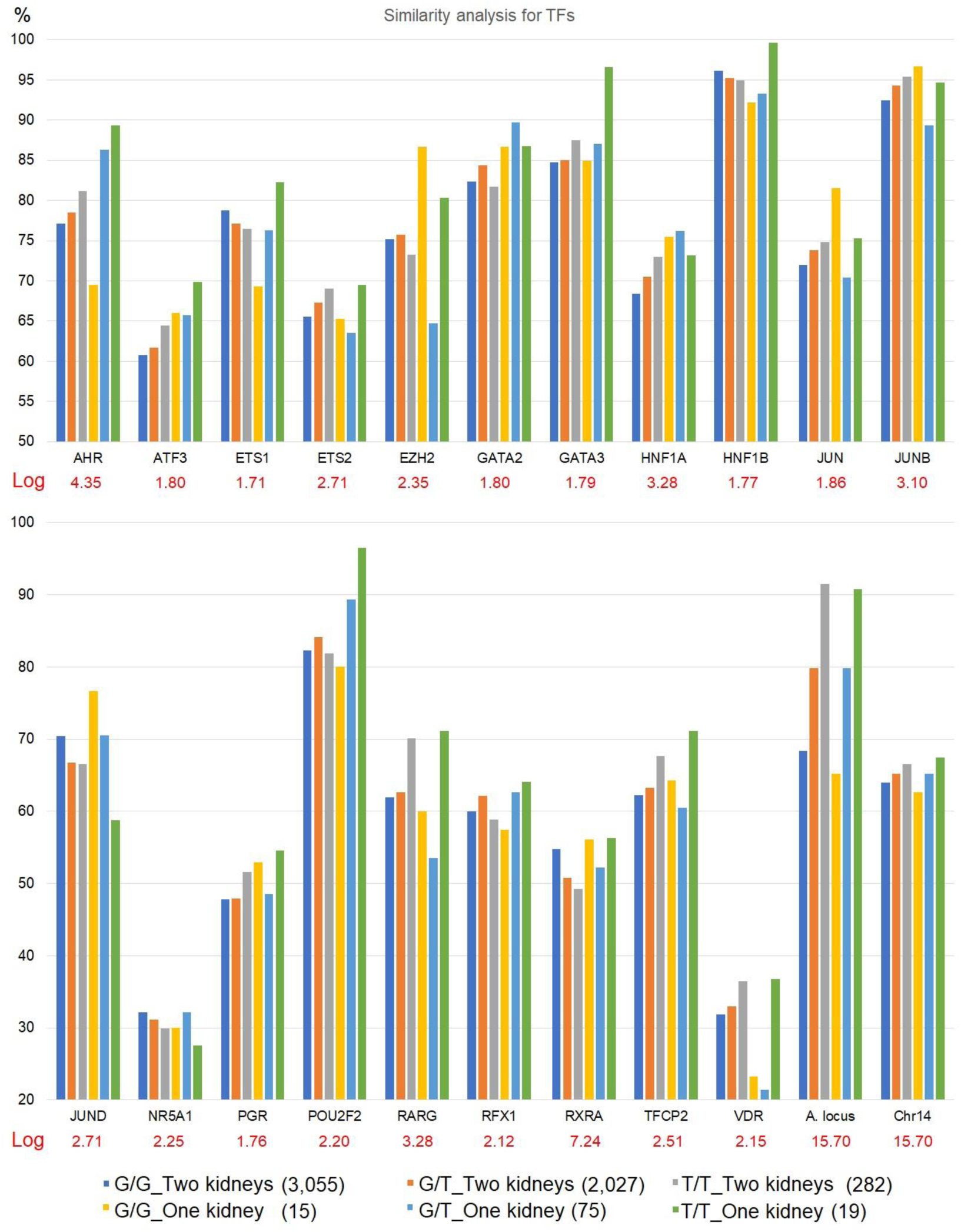
The genic similarity score for genes encoding TFs able to interact with the Erv insertion located on intron 1 of *KIT*. Animals were grouped based on genotype and phenotype interaction. Genotype grouping was performed using the most significantly associated SNP, chr14:36,411,266. The Jaccard index was used to construct a similarity index using the HSRA strain as the reference. Ten-percent FDR significance threshold for 107 TFs (tests) was applied (- log10 equal to 1.65). The A. locus represents the URA-associated locus chr14:32.9-36.6 Mb. The number of individuals included in each group was: 15 (G/G_One kidney), 3,055 (G/G_Two kidneys), 75 (G/T_One kidney), 2,027 (G/T_Two kidneys), 19 (T/T_One kidney) and 282 (T/T_Two kidneys). SEM values are not shown since they were estimated to be by order of 1 in 1,000 for all groups, given the high sample size (n = 5,576).

## 5. Discussion

Kidney organogenesis is the result of a complex cascade of processes, including budding, reciprocal inductive tissue interactions, stem cell growth and differentiation, cell polarization, mesenchymal to epithelial transformation, branching morphogenesis, angiogenesis, apoptosis, cell fusion, proximal-distal segmentation, and differentiation of multiple cell types (Schwab et al., 2003). URA is caused by the failure of embryonic kidney formation, a process initiated at the fifth gestational week in humans (Elumalai and Mampa, 2017).

Kamba et al. (2001) described a mechanism for URA in mice when analyzing FUBI (failure of ureteric bud invasion) mice embryos, performing an assessment through embryonic development. FUBI strain shows a URA incidence of 60% (imperfect penetrance). The outgrowing ureteric bud of the mesonephric duct interacts with the metanephric mesenchyme tissue. This interaction orchestrates embryonic kidney formation, which fails when the ureteric bud cannot develop at an early fetal growth stage, disrupting the branching morphogenesis process. Ureteric buds and the metanephric rudiments communicate in normal embryos. The ureteric buds develop branches on both sides in the renal rudiments through an invasive process. These branches end up as well-developed arborizations. Invasion of the metanephric mesenchyme on one side (less frequently on both sides) sometimes fails. The ureteric bud cannot contact and invade the metanephric mesenchyme. Instead of showing arborization, each ureteric bud has blind ends. At E12.5 in mice, the metanephric rudiment that did not experience invasion shows significant apoptosis and complete reabsorption by E13-E14. However, URA seems to be a multifactorial phenotype. For instance, not all cases result from incompetent ureteric bud development. Complete in utero involution of embryonic kidneys leading to URA has also been described (Elumalai and Mampa, 2017). This multifactorial phenotype classification might make genetic analysis more difficult since several pathways and mechanisms might result in the same phenotype after embryogenesis is accomplished. As a result, it could contribute to reducing power for gene detection.

### 5.1 Whole genome association analysis for unilateral kidney agenesis

Despite the potential existence of multiple mechanisms driving URA in rats, a highly significantly associated region showing the existence of a predominant URA mechanism in the present population was found. The most highly associated region on chromosome 14, 32.9 to 36.6 Mb, harbors several characterized genes (Figure 2). The two more numerous groups were lincRNAs and protein-coding genes. All the protein-coding genes identified are expressed in kidney-associated tissues and related structures (Supplementary Table 4). The associated locus harbors the KIT gene. The KIT gene was previously reported as the most likely candidate gene for URA in HS rats (Shull et al., 2006; Solberg Woods et al., 2010; Becker et al., 2015). *KIT* is involved in regulating cell proliferation, survival, and migration. Given this role, it is a candidate gene for multiple types of cancer (Hirota et al., 1998; Kitamura et al., 2001; Cho et al., 2006; Lück et al., 2010; Koelz et al., 2011; Donnenberg et al., 2012; Janostiak et al., 2018). It has oncogenic and tumor suppressor functions depending on tissue type (Janostiak et al., 2018), and it is used as a medical target to treat cancer. The *KIT* gene was also associated with coat color and deafness in cats. David et al. (2014) reported that a homologous polymorphism to the Erv insertion identified by Shull et al. (2006), Solberg Woods et al. (2010) and Becker et al. (2015) in rat, a 7,125 bp feline endogenous retrovirus (Ferv1) insertion as responsible for pigmentation variation and deafness. This insertion was associated with a pleiotropic effect. The Ferv1 insertion promotes complete penetrance for the absence of coat pigmentation and incomplete penetrance for deafness and iris hypopigmentation. This phenotypic effect was suggested as being caused by disruption of a DNAase I hypersensitive site in intron 1 of *KIT*.

The risk allele identified in HS rats for the leading associated SNP (chr14:36,411,266) was the allele T; however, some cases lack this allele, making it evident that more than one genetic mechanism associated with URA might be involved. It is also apparent that the risk allele is not enough to cause URA since only 6.49% of TT individuals expressed this phenotype, being URA elements in the genetic background probably involved and able to modify case presentation. The genomic similarity score distribution (Figure 7) supports this mechanism since T/T_One kidney is more similar to HSRA even after excluding the associated locus. Cases with genotype T/T showed the highest similarity to the HSRA strain. On the other hand, G/G cases are the least similar. Based on this result, it is possible to theorize that the founder family selected to generate the HSRA strain might be genetically closer to T/T cases. The assessed rat population two showed a high case-control imbalance and a low number of cases, including only nine individuals with URA (proportion 1:174), compromising the association analysis. The segment Chr14:33.8-34.1 Mb was associated with URA in the population three (Figure 3c), agreeing with the present analysis and previous reports, which also identified this chromosomal region as associated with URA (Shull et al., 2006; Solberg Woods et al., 2010; and Becker et al., 2015; Hansen and Spuhler, 1984). The population three had a lower marker density, explaining the presence of a non-completely overlapping associated region with the analyzed HS population.

*KIT* encodes a receptor involved in kidney organogenesis. It regulates the ureteric bud’s invasion of the metanephric rudiments, branching, and final arborization (Sanchez-Ferras et al., 2021). Sanchez et al. (2021) identified four major cell populations that serve as the progenitor for the nephric duct during morphogenesis, Nephric duct Progenitor 1-4 (NdPr1-4). Taguchi and Nishinakamura (2017) used KIT and CXCR4 proteins to isolate and describe these nephrotic duct progenitor cells; KIT was enriched in NdPr2 and NdPr4 cells. Schmidt-Ott et al. (2006) showed that in mice, histologically, KIT was abundant around the entry point of the ureteric bud into the metanephric mesenchyme rudiments. It was also strongly expressed in a multicellular layer of cells surrounding the entire metanephric mesenchyme at E11.5. The whole developing kidney is surrounded by KIT-positive cells by E13.5. The ureteric bud stalks are densely surrounded by KIT-positive cells before initiating an invasive process of the interstitium. This invasion process is performed by branches developed from the ureteric bud. Additionally, S-shaped bodies expressing *KIT*, start being identifiable before noting initiation of KIT expression repression.

Arnould et al. (2009) identified *KIT* as key for kidney development and MMP9 as the protein able to perform lysis of the KIT receptor, releasing its active form, the stem cell factor (SCF). Arnould et al. (2009) and Bengatta et al. (2009) reported that MMP9 deficiency impeded embryonic kidney maturation through the generation of less SCF, which in turn increases apoptosis (between 2.5 to 5-fold). This delayed maturation was associated with branching defects, 30% fewer nephrons, 20% lighter kidneys, and abnormal architecture at 12 months. Schmidt-Ott et al. (2006) reported that inhibition of *KIT* generated quantitative reductions in ureteric bud branching, the number of glomeruli, ureteric bud tips, and total nephrons.

### 5.2 Polygenic epistasis assessment and founder haplotype mosaic estimation

The association analysis performed using GxTheta showed a low number of polymorphisms whose effect is modified by the genetic background for specific strains (one polymorphism for ACI, two for BN, and two for BUF). GxTheta tests “polygenic epistasis.” It analyzes if an allele’s effect depends on the global genetic ancestry proportion, in this case, from the eight original founder strains. As a result, the effect of the tested polymorphisms can be assumed to be same across all eight genetic backgrounds (figure 4). The examined polymorphisms do not show widespread epistasis with multiple loci across the genome (Rau et al., 2020). Figure 5 and Supplementary figure 1 show the estimation for haplotype composition in cases and controls. For the most significantly associated region, chr14:32.9-36.6 Mb, ACI influences majorly the presentation of URA; however, the risk allele and the top one hundred associated polymorphisms are not ACI exclusive (figure 6a); the same alleles are present in ACI, BUF and M520 (upstream-35Mb). URA incidence in BUF and M520 is lower than in ACI, and these alleles do not depend on the global genetic ancestry proportion, suggesting that their effect is equivalent between ACI, BUF, and M520. Another highly significant locus in cases showing a higher probability of being inherited from ACI, the downstream-35Mb, harbors several ACI exclusive polymorphisms (light blue dots in figure 6a). This locus was not identified as showing polygenic epistasis since the effect of these alleles cannot be tested for the remaining strains. The downstream-35Mb might be involved in modifying URA penetrance.

### 5.3 HSRA similarity score analysis

The similarity analysis aimed to identify potential loci able to modify penetrance for URA. For the HSRA strain, the number of URA elements were increased through phenotypic selection, causing a rise in URA incidence from around 1.9% (incidence in the present HS population and probably for the original HS population source of the HSRA strain) up to 75%. The genomic similarity score shows that T/T animals (for chr14:36,411,266) are slightly (0.93%) more similar to HSRA than G/G individuals (Figure 7). However, when analyzing only the URA-associated region chr14:32.9-36.6 Mb (Figure 8), this difference is more evident, going from 65.2% for G/G_One kidney up to 90.8% for T/T_One kidney. This result is related to the limitations of the similarity score analysis *per se*. This analysis depends on the number of elements used for calculation since it is an average value across polymorphism by analyzed locus. The G/G vs. T/T difference also supports the theory that founder HSRA individuals might have been genetically closer to T/T than G/G genetic architecture. This results suggests the existence of at least two URA mechanisms (T/T and G/G at chr14:36,411,266) in this HS population. In HSRA, the higher similarity for T/T animals shows that this locus was further selected. Since similarity for this locus is lower than one hundred, HSRA individuals show an additional selection pressure on the associated locus driven by phenotypic selection. Urmo et al. (2021) and Munro et al. (2022) reported the presence of cis-eQTLs for *KIT* in this HS population in the brain.

The associated locus chr14:32.9-36.6 Mb fails to differentiate always phenotypic status, implying that other regions different from the associated locus might be involved in modifying URA incidence in the present population. The similarity analysis for TFs able to interact with the Erv insertion in intron 1 of KIT (figure 8) was used to identify loci able to modify penetrance. Some loci showed the highest HSRA similarity score for T/T_One kidney, including Aryl Hydrocarbon Receptor (*AHR*), Activating Transcription Factor 3 (*ATF3*), ETS Proto-Oncogene 1, Transcription Factor (*ETS1*), ETS Proto-Oncogene 2, Transcription Factor (*ETS2*), GATA Binding Protein 3 (*GATA3*), HNF1 Homeobox B (*HNF1B*), Progesterone Receptor (*PGR*), POU Class 2 Homeobox 2 (*POU2F2*), Retinoic Acid Receptor Gamma (*RARG*), Regulatory Factor X1 (*RFX1*), and Transcription Factor CP2 (*TFCP2*).

Some of the identified TFs are described as critical for kidney organogenesis, including *AHR*, *ETS1*, *GATA3*, *HNF1B,* and *RARG*. During nephrogenesis, *AHR* modulates mesenchymal-to-epithelial transition through the regulation of *WT1* (Ramos, 2006). Falahatpisheh and Ramos (2003) and Ramos (2006) reported that unregulated activation of AHR signaling represses glomerulogenesis and branching morphogenesis of metanephric kidneys; it also decreases comma- and S-shaped bodies, numbers of glomeruli, and tubulo-epithelial structures. Activation of AHR stalls the differentiation of glomerular cells. These processes require alternative splicing of *WT1* and prevent glomerulogenesis and tubulogenesis. AHR stalls podocyte differentiation and promotes several hyperproliferative phenotypes. Ureteric bud development was affected by AHR activation. Dysregulated AHR signaling promoted a significantly lower number of ureteric bud branching points in metanephric cultures. Decreased branching was present at second and third tier branching points (Ramos, 2006).

In mice, disruption of *GATA3* generates bilateral renal agenesis coupled with deficient nephrotic duct elongation and severe renal hypoplasia (Grote et al., 2006; Sanchez-Ferras et al., 2021). Nephric duct Progenitor 1 to 4 (NdPr1-4) cells used by Sanchez-Ferras (2021) surge in time and space in a stereotypical pattern, and progenitor cell progression is tightly regulated by TFAP2A/B and GATA3 TFs; *GATA3* expression increases from the rostral portion of the nephric duct to the caudal (tip) of the same structure. Deactivated *GATA3* was coupled with nephric duct elongation defects, resulting in a massive increase in nephric duct cellularity and aberrant elongation of the nephric duct. These defects generate kidney agenesis (Grote et al., 2006; Sanchez-Ferras et al., 2021). Knockout *GATA3* animals revealed a 40% decrease in elongation at E9.5; this structure is aberrantly shaped at E10.5, showing a pronounced number of cells per duct section and enhanced lumen size. Knockout animals also show accumulation of NdPr1 and NdPr2 identity and no progression towards NdPr4, being NdPr4 the precursor of the ureteric bud.

Regulation of KIT by TFs showing selection involving a trans-eQTLs effect was previously reported. Urmo et al. (2021) identified the presence of trans-eQTLs for *KIT* using human blood. The *HNF1B* TF was reported as having a *KIT* trans-eQTL (Table 6). This *KIT*-HNF1B trans-eQTL confirms potential additional elements for URA able to explain incomplete penetrance in the present analysis. *HNF1B* has expression in the Wolffian duct and ureteric bud epithelia; this gene regulates ureteric arborization formation, collective duct differentiation, pronephros size, initiation of nephrogenesis, nephron segmentation, and proper tissue architecture maintenance (Sauert et al., 2012; Ferrè and Igarashi, 2019). *HNF1B* regulates ureteric arborization formation, collective duct differentiation, and tissue architecture. Desgrange et al. (2017) reported that *HNF1B* regulates cell-cell contacts and apicobasal polarity during early branching events. Dysregulation of HNF1B generates severe epithelial disorganization and lower cell reorganization during mitosis-associated cell dispersal. Defects on this TF also stall critical players of the GDNF-RET pathway (Sauert et al., 2012). Therefore, HNF1B is crucial during kidney organogenesis. Sauert et al. (2012) suggest that activity of HNF1B is performed by regulating genes responsible for processes such as transport and intrinsic cell-membrane components. Paces-Fessy et al. (2012) reported that kidney hypoplasia, caudal ectopic aborted ureter buds, duplicated kidneys, megaureters, and hydronephrosis are phenotypes associated with heterozygous HNF1B-/+ PAX2-/+ individuals progeny from knock-out (KO) subjects. These animals also showed delayed nephron segmentation, medullar interstitial differentiation, increased apoptosis, and transitory *LIM1*(*LIM Homeobox 1*) and *WNT4* (*Wnt Family Member 4*) downregulation. Niborski et al. (2021) and Oram et al. (2010) identified *HNF1B* as generating simultaneous genital tract anomalies. However, it rarely shows an association with isolated uterine abnormalities.

**Table 6.**
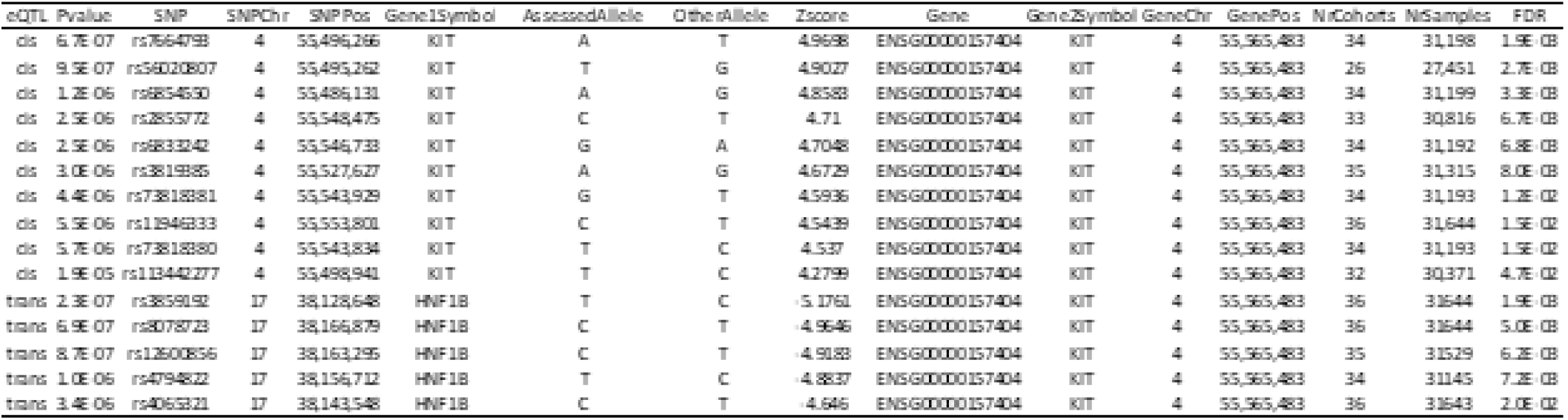
Human cis and trans-eQTLs involving *KIT* identified by Urmo et al. (2021) in blood. The genomic location of the human *HNF1B* is chr17:37,6-37,7 Mb.

Using KO *ETS1* mice, Ye et al. (2018) identified an overrepresentation of structural kidney defects, including unilateral duplicated ureters, duplicated kidneys, and unilateral renal hypoplasia. However, KO mice do not show complete dominance. ETV4 and ETV5 belong to this family, positively regulated by RET signaling in the ureteric bud tips. RET is a crucial receptor tyrosine kinase involved in the branching morphogenesis of the ureteric bud (Lu et al., 2010). Lu et al. (2010) also showed that animals with a double KO allele for ETV4 and one ETV5 allele display either severe hypodysplasia or renal agenesis; however, double KOs show kidney agenesis. Retinoic acid receptors, a family of TFs, are crucial for controlling RET expression in the ureteric bud. Dominant-negative Retinoic acid receptor stalls RET expression and its pathway signaling, repressing ureteric bud formation and branching morphogenesis (Rosselot et al., 2010). Rosselot et al. (2010) reported that in the embryonic kidney, RET expression and branching depend on RALDH2. Batourina et al., (2001) reported that induced RET expression, a maturation ureteric bud marker (Taguchi and Nishinakamura, 2017), in double KO Retinoic Acid Receptor Alpha (RARA) and Retinoic Acid Receptor Beta (RARB2) restores ureteric bud growth.

### 5.4 A possible mechanism for URA in HS

It is hypothesized that the mechanism underlying URA identified in the present rat population might be similar to the one described by Kamba et al. (2001) in FUBI mice. This mouse strain shows a failure of ureteric bud invasion. The ureteric bud is the defective structure. There is asymmetric ureteric bud branching failure. The metanephric rudiment initiates an apoptotic process since there is no ureteric bud invasion, generating complete structure absorption. This process being mediated by TFs (Figure 9) able to interact with sequences of the Erv insertion in intron1 of KIT (Figure 6c), a gene mediator of the ureteric bud formation, progression, and invasion of the ureteric bud through the metanephric rudiments on both sides, a key process for kidney organogenesis. This theory is also supported by the identification of TFs such as *HNF1B* as being potentially responsible for increasing penetrance since *HNF1B* regulates ureteric arborization formation, collective duct differentiation, and tissue architecture. Therefore, unilateral ureteric bud invasion might be the impaired kidney organogenesis step being stalled in HS rats.

**Figure 9.**
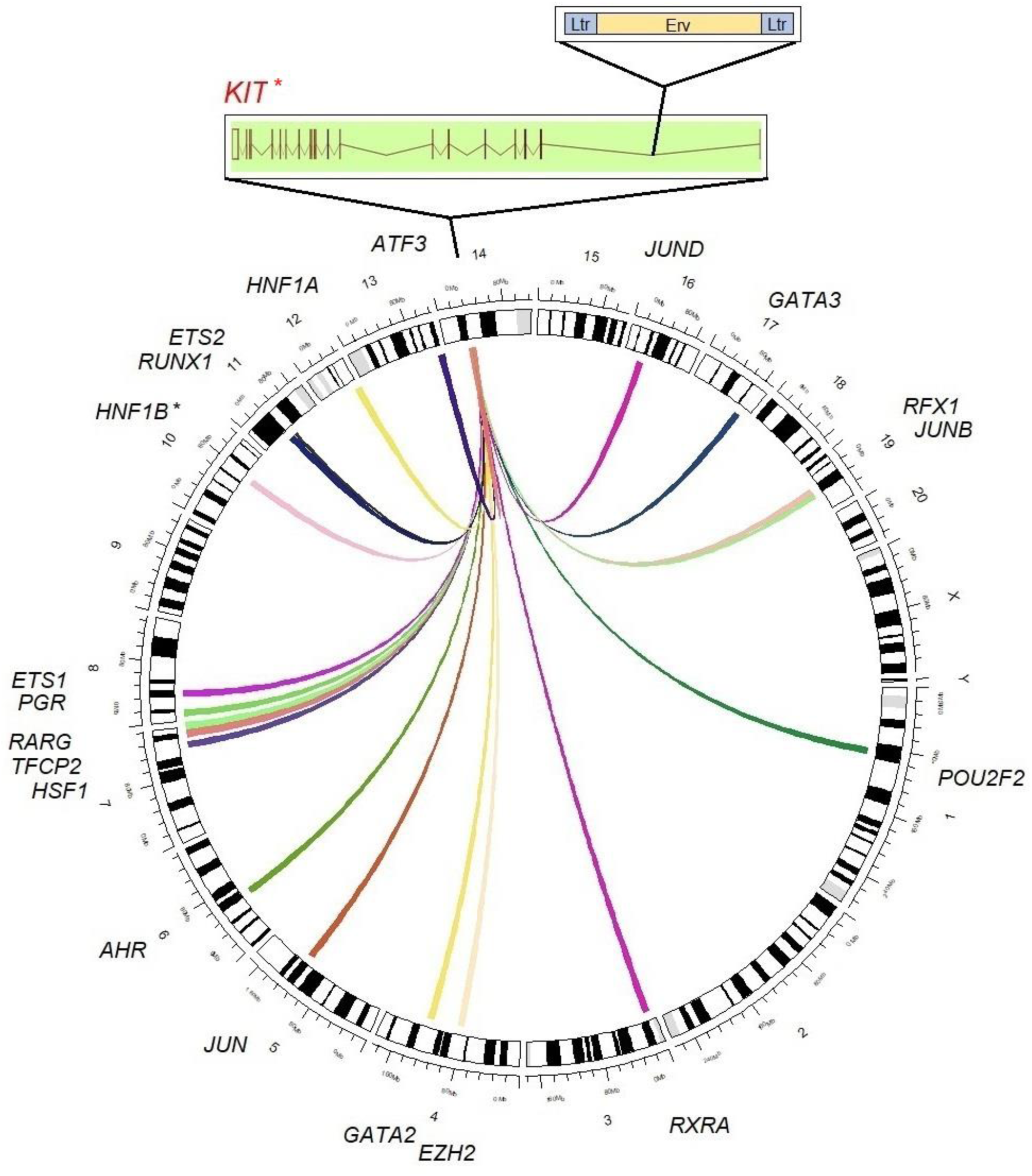
Rat ideogram for potential URA-related elements in HS rats. Loci showing evidence of selection in HSRA and higher similarity for T/T_One kidney which were selected as candidates for increasing URA penetrance in HS rats. *KIT* cis-regulated related elements are represented by the salmon link on *KIT*. *KIT* trans regulators are represented by additional links on the *KIT* locus. The location of the Erv insertion inside the *KIT* gene is presented. *Cis and trans eQTLs reported by Urmo et al. (2021) in human and by Munro et al. (2022) in HS rats.

The similarity analysis for TFs able to interact with the Erv insertion in intron 1 of *KIT* shows selective pressure in the reference strain HSRA. Some loci showed high HSRA similarity scores for T/T_One kidney, including *AHR*, *ATF3*, *ETS1*, *ETS2*, *GATA3*, *HNF1B*, *HSF1*, *PGR*, *POU2F2*, *RARG*, *RFX1*, and *TFCP2*. Some of these TFs are reported as involved in kidney organogenesis. Additionally, to general cellular architecture, several steps in the organogenesis cascade are paramount. Some of these steps are reported as modulated by these TFs, including the number of ureteric bud tips, branches and branch ramifications, total branch elongation, and lumen size. All these parameters are indicators of KIT activity and modulate ureteric bud invasion of the metanephric rudiment; therefore, they can regulate the final number of nephrons in the mature kidney. These features have a quantitative description, and URA might result from a cumulative additive defect generated by KIT activity and *KIT* regulator TFs. For this reason, these loci might contribute to the imperfect penetrance of URA in HS rats. This mechanism implies the existence of a minimum threshold for the final number of nephrons required for stalling the apoptotic process of the metanephric rudiments. A minimum number of ureteric bud tips, branches, and branch ramifications, total branch elongation, and lumen size could be required to continue the organogenic molecular cascade up to a mature kidney, which might be the basis for imperfect penetrance for kidney agenesis in HS rats. However, the existence of alternative organogenesis pathways and involved genes needs to be considered.

## 6. Conclusion

A URA-associated region on chromosome 14, 32.9 to 36.6 Mb, harboring the *KIT* gene was found. This gene was previously reported as the most likely candidate for URA in HS rats, a gene involved in cell proliferation, survival, and migration regulation. An Erv insertion was found inside the intron one of the *KIT* gene, and divergent insertion composition was identified for HS founder strains, and the URA selected strain, HSRA. This Erv insertion was previously reported as the most likely candidate polymorphism responsible for this congenital urinary tract malformation. Given the presentation of low penetrance for this phenotype in HS rats, a similarity analysis aiming at identifying potential loci able to modify penetrance for URA was performed using sequences from elements of the identified Erv insertion. Applying this methodology, several TFs able to interact with the Erv insertion show selection in HSRA and higher similarity with T/T_One kidney rats were identified, including *AHR*, *ATF3*, *ETS1*, *ETS2*, *GATA3*, *HNF1B*, *HSF1*, *PGR*, *POU2F2*, *RARG*, *RFX1*, and *TFCP2*. These TFs were recognized as potential candidates responsible for increasing URA penetrance. A mechanism categorizing URA as a threshold phenotype was suggested in HS rats. It implies the existence of a minimum threshold for the final number of nephrons required for stalling the apoptotic process of the metanephric rudiments, a minimum number of several structures, including ureteric bud tips, branches, branch ramifications, total branch elongation, and lumen size. A minimum number of functional kidney-associated structures could be required to continue the organogenic molecular cascade up to a mature kidney, which might be the basis for imperfect penetrance recognized for kidney agenesis. Individuals with this quantitative cumulative defect would exhibit URA with decreased penetrance.

## 7 Acknowledgments

We would like to thank Dr. Takashi Kuramoto from the Department of Animal Sciences, Tokyo University of Agriculture for providing information about their *KIT* Erv insertion.

